# State-dependency of behavioural traits is a function of the life-stage in a holometabolous insect

**DOI:** 10.1101/2022.12.15.520519

**Authors:** Pragya Singh, Jonas Wolthaus, Holger Schielzeth, Caroline Müller

## Abstract

State variables, such as body condition, are important predictors of behavioural traits. Depending on the state of an individual, the costs and benefits associated with different behavioural decisions can vary. An individual’s state could affect its average behavioural response and also the behavioural repeatability. Moreover, even for the same state, different behavioural strategies may be adaptive depending on the individual’s life-stage. Here, we manipulated the body condition in larvae and adults of a holometabolous insect species, *Athalia rosae*, using starvation. We assessed the effects of starvation on the behavioural traits of post-contact immobility (PCI) and activity levels and tested their repeatability. Our results show state-dependency of behaviour, although the effect varied with life-stage. Starved larvae exhibited shorter PCI duration and higher activity levels, whereas starved adults were less active than non-starved individuals. Moreover, although most behavioural repeatability estimates were significant in both life-stages, we did not find any significant effect of starvation on the estimates. Next, we calculated standardised effect sizes to compare starvation effects across life stages. We found that starvation had a larger and opposite effect in the larval stage than during the adult stage for all behavioural traits. Finally, we conducted microcosm and no-choice bioassay experiments to examine the benefits and costs, respectively, of the behaviour elicited by starvation in the larval stage. We observed that starved larvae located food faster than non-starved larvae but were also attacked sooner by a predator, possibly due to their higher activity levels. Together, our results demonstrate that behavioural state-dependence is a function of the life-stage of an individual. Moreover, the behavioural strategy exhibited can be adaptive for a specific life-stage with respect to certain functions, like foraging, but also carry costs, like risk of predation.

## Introduction

State variables, such as body condition or nutritional status, are often important predictors of individual-specific behaviour (Sih *et al*., 2015). Depending on the state of an individual, the costs and benefits associated with different behavioural decisions can vary (Houston & McNamara, 1999; Dingemanse & Wolf, 2010; Luttbeg & Sih, 2010). This variation between individuals in state can then translate into between-individual differences in behavioural traits (Niemelä & Dingemanse, 2018; Edmunds *et al*., 2021). Moreover, the life-stage at which an individual experiences a state could be an important determining factor for the expressed behaviour (Sherratt & Morand-Ferron, 2018; Moran *et al*., 2021; Rádai *et al*., 2022). Even for the same state, different behavioural strategies may be adaptive depending on the life-stage at which the state is experienced by an individual. This is particularly important for organisms that have complex life-cycles or experience ontogenetic niche shifts (Wilson & Krause, 2012; Stamps & Krishnan, 2017; English & Barreaux, 2020; Cabrera *et al*., 2021).

Several models have been proposed regarding state-dependent behaviour. Some theoretical models predict that individuals in poor body conditions (e.g. with lower energy reserves) will exhibit more aggression, boldness or activity levels because they have a greater ‘need’ of food and have relatively less to lose (Rands *et al*., 2003; Dall *et al*., 2004), while individuals in good conditions may be more cautious, i.e. shyer and less active, to protect what they have (the ‘asset-protection principle’, Ludwig & Rowe, 1990; Clark, 1994). These are considered ‘needs-based’ explanations (Barclay *et al*., 2018). In contrast, individuals in good body condition (e.g. with higher energy reserves) may also behave more active and bolder if their better nutritional state provides them with a higher survival chance even in risky situations, e.g. by being better at escaping predators, as postulated in the ‘state-dependent safety hypothesis’ (Luttbeg & Sih, 2010; Moran *et al*., 2021). This offers an ‘ability-based’ explanation (Barclay *et al*., 2018). How valid these different models are, may depend on the life-stage, because selection pressures vary across ontogeny, with growth being important for early developmental stages and reproduction for adults, leading to stage-specific behavioural strategies. While an individual’s state could affect its average behavioural response, it could also impact the behavioural repeatability or consistency (Luttbeg & Sih, 2010; Sih *et al*., 2015; MacGregor *et al*., 2021 but also see Niemelä & Dingemanse, 2018). Furthermore, there can be state-dependant feedback loops that can affect individual behavioural differences (Luttbeg & Sih, 2010; Sih *et al*., 2015), which in turn can impact ecological processes (Sih *et al*., 2012). Usually, stressful conditions increase phenotypic variation and plasticity, thus we may expect a decrease in behavioural repeatability under stress.

One way of manipulating an individual’s body condition or state is by exposing it to starvation, which usually depletes energy reserves (Arrese & Soulages, 2010). Starvation or food limitation is a ubiquitous stress faced by organisms in the wild with food availability varying temporally or/and spatially, and leading to effects on behaviour (Scharf, 2016; Müller & Müller, 2017), life-history traits (Singh *et al*., 2020; Paul *et al*., 2022), metabolism (Zhang *et al*., 2019) etc. For example, the behavioural activity level can change in response to starvation, with some species showing an increased activity, while others showed a decrease and few also showed a hump-shaped response (i.e. an increase followed by a decrease) (Scharf, 2016). However, how the response to starvation in activity levels changes across life-stages within the same organism is less-well studied.

Here, we investigate the effects of starvation on behavioural traits across ontogeny, i.e. in the larval and adult life-stage, using the turnip sawfly, *Athalia rosae* (Hymenoptera: Tenthredinidae). Sawfly larvae feed on various Brassicaceae plants and pupate in the soil. The adults feed on floral nectar, usually from Apiaceae plants. During the larval stage, individuals can rapidly defoliate their host plants and hence experience periods of starvation (Riggert, 1939). Likewise, eclosing adults may not immediately have access to nectar, leading to periods of starvation. In the present study, we investigate post-contact immobility (PCI) (Sendova-Franks *et al*., 2020) and activity levels in larvae and adults. PCI is a behavioural strategy following physical contact with the predator, in which the individual is immobile for some duration. PCI is also known as post-predation immobility, tonic immobility, thanatosis or ‘death-feigning’ behaviour. PCI is proposed to function as an anti-predator strategy by increasing survival chances (Rogers & Simpson, 2014; Humphreys & Ruxton, 2018; Skelhorn, 2018; Farji-Brener *et al*., 2022), and has often been used as a proxy for boldness or risk-taking behaviour in animals (Edelaar *et al*., 2012; Tremmel & Müller, 2013; Segovia *et al*., 2019). Furthermore, we examined general activity, measured as the distance moved and time spent immobile (immobility duration) during a defined observation period. The chosen behavioural traits of PCI and activity reveal ecologically relevant information about individuals regardless of their ontogenetic life-stage and niche (Wilson & Krause, 2012).

First, we investigated in laboratory assays whether (i) the PCI duration, and (ii) activity levels varied in response to starvation duration in the larval or adult stage and assessed the effect of starvation on behavioural repeatability within these stages. Next, we calculated standardised starvation effect sizes for larvae and adults to facilitate life-stage comparisons. Since the starvation effect was larger during the larval stage, we examined the potential benefits and costs of the behavioural strategy induced by larval starvation. For this, we assessed the effect of larval starvation on (i) locating food, and on (ii) latency of attack by a potential predator using microcosm and bioassay experiments, respectively. We hypothesised that if the ‘asset-protection principle’ holds in our study system, starvation will lead to increased activity levels and reduced PCI duration, while we would expect the reverse if the ‘state-dependent safety hypothesis’ holds. Moreover, we expected that larvae would be affected more acutely by starvation than adults, as foraging is the crucial activity for larvae. We also expected starved individuals to show lower repeatability than control individuals. Lastly, we expected increased activity levels due to larval starvation to lead to increased foraging success, but also to have costs in terms of increased predation. By addressing these hypotheses, our study increases our understanding of ontogenetic effects on state-dependency of behaviour, and the adaptive value of different behavioural strategies across life-stages in a holometabolous insect.

## Materials and Methods

### Experimental Sawflies

Sawflies for this study were derived from a laboratory stock population, which has been established from *A. rosae* adults collected in the surroundings of Bielefeld, Germany. The stock is supplemented annually with field-caught insects. The sawflies were maintained in mesh cages (60 × 60 × 60 cm) in a laboratory with a 16 h: 8 h light: dark cycle, approximately 60% relative humidity and at room temperature. Several adult females and males were released in a cage with plants of *Sinapis alba* (Brassicaceae) for oviposition, leading to both unfertilised and fertilised eggs, resulting in male and female offspring, respectively, as *A. rosae* is haplodiploid. After 6-7 d, the emerging larvae were reared on non-flowering plants of *Brassica rapa* var. *pekinensis* (Brassicaceae). The *S. alba* and *B. rapa* plants were grown from seeds in a climate chamber (20 °C, 16 h: 8 h light: dark, 70% r.h.) and greenhouse (≥ 20 °C, 16 h: 8 h light: dark, 70% r.h.), respectively. There were minor variations in cage set-up for different experiments as detailed below.

### Larval behavioural traits

#### (a) PCI assay

Approximately 40 unmated females were put in a single cage for 24 h for oviposition. Females were then transferred to another cage, and allowed to oviposit for another 24 h. Only unmated females were used in this experiment to ensure that all eggs laid are unfertilised and hence, all offspring larvae are males, as there can be PCI differences between sex (Miyatake, 2001). Twenty days after cage set-up, larvae (4^th^-5^th^ instar) were collected from each cage and put individually in Petri dishes (5.5 cm diameter) lined with moist filter paper (moistened with tap water). Each larva was weighed initially and randomly assigned to one treatment, control and starved, and again weighed at the end of the treatment after 4 h (Sartorius ME36S Balance) to confirm that starved larvae lost more body mass than control larvae (Supplement S1). In the starvation treatment, the larvae had no access to *B. rapa*, while the control treatment larvae were provided *B. rapa* leaf discs ad-libitum over the 4 h period. To induce PCI, a larva was grasped from the middle of the body using forceps and dropped from a height of 2 cm onto a Petri dish. This handling using forceps was used to mimic a predation attack and induce PCI. If in the first trial the larva did not show PCI, the forceps stimulation was repeated once or twice until a PCI was recorded. A larva was considered as exhibiting PCI if it curled up its body tightly, and stayed immobile in this posture for at least a second. The PCI duration was measured from the time the larva entered PCI posture until it uncurled its body completely and moved at least 1 mm. Each larva was observed for a maximum of ten minutes. The PCI duration was measured repeatedly once every hour over a period of 4 h. The sample size for larvae of the starvation and control treatment were 25 and 24, respectively.

#### (b) Activity levels assay

Adult female and male sawflies were set-up in a single cage for 24 h for oviposition. Twenty days later, we collected thirty larvae (4^th^-5^th^ instar) and randomly assigned them to a starvation or control treatment (n=15 for each treatment). Larvae were weighed before the experiment and put in individual Petri dishes lined with moist filter paper. Similar to above, starvation treatment larvae had no access to *B. rapa*, while the control treatment larvae were provided *B. rapa* leaf discs ad-libitum. The activity level of each larva was measured after 2 h and 4 h of initiating the treatment. For measuring their activity levels, larvae were moved individually to empty Petri dishes, and their behaviour was recorded and tracked for 10 min using Noldus Ethovision V7 (Wageningen, NL; centre-point tracked at 1.92 samples/s). We recorded distance moved and immobility duration for each individual. Six individuals were recorded simultaneously in one trial using a single overhead camera. We limited the observation period to ten minutes as a longer duration could have led to starvation effects in larvae of the control treatment.

### Adult behavioural traits: PCI and activity levels assay

Freshly eclosed adults were collected from the laboratory stock population daily over a period of multiple days. We determined the sex of the adults and randomly assigned them to a starvation or control treatment. Each individual was weighed (Sartorius CPA224S-OCE Balance) and then put in an individual Petri dish lined with moist filter paper. Starved and control individuals were provided water only or a 2% (v/v) honey-water solution, respectively, on tissue-paper, which was replenished every other day. Two and 4 d after eclosion, we measured the PCI duration and activity levels of each individual. For measuring the PCI duration, an individual was grasped from the middle of the body using forceps, held for five seconds and then placed in the Petri dish head first. An adult was considered as exhibiting PCI, if it curled up and remained immobile for at least one second. Each adult was observed for a maximum of eight minutes. The PCI duration was counted until the adult stood upright on its legs. Thirty minutes after initiating the PCI assay, individuals were put in empty Petri dishes and their behaviour was recorded and tracked using Noldus Ethovision V7 (settings similar to above) for 1 h. We recorded distance moved and immobility duration for each individual. For females, the sample size was 30 for each treatment for PCI and activity parameters. For males, the PCI sample size for starvation and control treatment were 29 and 27, respectively. For the activity parameters, the sample size was 27 males for each treatment as we lost two individuals that escaped after the second PCI measurement.

### Microcosm experiment to examine effect of larval starvation on locating food

Larvae (~14 days old, 4^th^-5^th^ instar) were collected from the cage set-up for the larval activity levels assay and randomly assigned to the starvation or control treatment, performed as described above (PCI assay). Each larva was kept individually in a Petri dish lined with moist filter paper for the duration. A cage (50 x 50 x 50 cm^3^) was set up with eight Petri dishes placed at equidistant locations from the centre. Each Petri dish had a *B. rapa* leaf disc (2.5 cm ⌀) and was covered with white paper to prevent visual detection. Eight larvae of the same treatment were put in a Petri dish without filter paper or leaves, and placed in the centre of the cage, and this was considered as one trial. Each trial lasted for 15 minutes. The timing to reach a leaf for a larva (if it did), and the number of leaves occupied by larva/larvae at the end of each trial were noted. Eight trials were conducted, with the larval treatment alternated with every trial (4 trials per treatment). New leaf discs were placed in the Petri dishes after every trial. To decrease directionality of olfactory cues, that larvae may use to locate leaves, we placed four *B. rapa* plants around the cage.

### No-choice bioassays to assess effect of larval starvation on predation

We used eleven fourth-instar individuals (length ~ 4 cm) of *Hierodula membranacea* (Mantidae) as potential predators (purchased from www.mantidenundmehr.de). The mantids were reared in individual cages (8 cm × 8 cm× 6 cm) in a climate chamber (approx. 25 °C) on a diet of *Drosophila melanogaster* and/or *Drosophila hydei*, and had not been exposed to *A. rosae* before the experiment. All mantids were starved for 48 h before the experiment. For the experiment, we collected similar-sized larvae of *A. rosae* from a laboratory culture cage (setup with male and female adults), and randomly assigned them to either starvation or control treatment, performed as described above (PCI assay). Each larva was kept individually in a Petri dish lined with moist filter paper for the treatment duration. After 4 h of treatment, we measured each larva’s body mass (range: 41.1-75.5 mg). Then, the no-choice bioassay was performed in a container (20 cm × 20 cm× 20 cm) that was covered with white paper from all sides to avoid any visual distractions to the mantid and larva. First the larva and then the mantid were added to the container at fixed positions facing each other. The larva was added to the container in a Petri dish cover, while the mantid was added on a meshed lid to the container. This allowed us to position the mantid ensuring that it faced the *A. rosae* larva without touching both individuals. If the larva showed PCI behaviour and curled up after being put in the container, we waited until the larva had uncurled before we added the mantid. We recorded the latency to attack by the mantid. During mantid attacks, we noticed that larvae showed easy bleeding (Müller & Brakefield, 2003), in which larvae exude haemolymph under mechanical stress, acting as a potential anti-predator mechanism (Boevé & Müller, 2005). After the attack, we removed the larva from the mantid’s grasp (if it had not dropped the larva itself) gently using forceps to ensure that the mantid remained hungry and to avoid any repercussions of consuming larvae, which contain glucosinolates sequestered from their host plants in their haemolymph (Müller *et al*., 2001). As the mantid would often be wiping its mouth to remove the haemolymph (similar behaviour described for lizards after attacking *A. rosae* larvae in Vlieger *et al*., 2004), it was easy to remove the larva. All assayed mantids were exposed to both treatment larvae alternatively, but in different treatment order of either starvation or control treatment first. Data from one mantid, which had recently molted, was excluded as it did not attack any larva.

### Statistical Analyses

All analyses were done in R version 4.0.4 (2021-02-15) (R Core Team, 2021). We included body mass for each analysis because body mass or size can have an effect on behavioural traits (Hozumi & Miyatake, 2005; Farkas, 2016). Variables were appropriately transformed to improve the distribution of model residuals. For the larval PCI assay, the effect of the treatment (Starvation or Control), time (1,2,3 or 4 h) and their interaction on log-transformed PCI duration were assessed using linear mixed models (lmm) (package ‘lmerTest’ version 3.1-3, package ’lme4’ version 1.1-30) (Bates *et al*., 2015; Kuznetsova *et al*., 2017), with body mass as covariate and individual identity as a random effect. As some PCI duration data points were zero, i.e. for individuals that did not show PCI, we added 1 to all data points before log-transformation to avoid infinite values. Note that all conclusions remained unchanged if cage was included as a factor (data not shown). For larval activity level assay, we examined the effect of treatment, time (2 or 4 h) and their interaction on distance moved and immobility duration using lmm, with body mass as covariate and individual identity as well as trial identity as random effects. For adult behavioural traits, we examined the effects of sex, treatment, time (2 or 4 day) and the interaction between treatment and time on PCI duration, distance moved and immobility duration using lmm, with body mass as covariate and individual identity as a random effect. Trial identity was also included as random effect for distance moved and immobility duration. For the larval microcosm experiment, we examined whether treatment had an effect on whether any leaf was reached using a binomial generalised linear mixed-effects model (glmm), with trial identity as random effect. We did not analyse the timing to reach a leaf or number of leaves occupied as very few control larvae had reached any leaf at the end of the trial (see below and S4). For the larval no-choice predation bioassay, we examined the effect of treatment on latency to attack (log-transformed) using a lmm, with larval body mass as covariate and mantid identity as well as treatment order as random effects. When interaction terms were non-significant, we dropped them to test the significance of the lower-order terms. The posthoc analyses were conducted using the package ‘multcomp’ version 1.4-17 (Hothorn *et al*., 2008).

#### Effect of starvation on behavioural consistency of individuals

We calculated the adjusted repeatability estimates (with time as fixed effect) (Nakagawa & Schielzeth, 2010) and associated confidence intervals (95% using parametric bootstrapping, n=1000) for the behavioural traits (PCI duration, distance moved and immobility duration) separately for control and starved individuals for larvae, adult males and adult females (’rptR’ package version 0.9.22, Stoffel *et al*., 2017). Likelihood ratio tests were used for significance testing of repeatability estimates. The traits were transformed as in the statistical analysis described above. The repeatability estimates allow us to quantify consistency in individual behaviour using individual identity for partitioning observed variance. The parametric bootstrapping of the ’rptR’ package simulates new data using the estimates from the original model and experimental design, and then calculates the repeatability of the simulated data. In that way, we obtained 1000 repeatability estimates for both starved and control treatment for each behavioural trait. To test if the repeatability estimates differed significantly between starved and control treatment for each trait, we calculated the difference between the repeatability estimates of starved and control treatments (subtracting the smaller from the larger estimate) based on parametric bootstrapping samples. We then examined what proportion of such differences was lower than zero, and calculated the asymptotic two-tailed *p*-value as twice this proportion.

#### Effects sizes

To facilitate life-stage comparisons of starvation effect, we calculated the standardised effect size Glass’ delta and its confidence intervals using the ’effectsize’ package v. 0.4.5 (Ben-Shachar *et al*., 2020). The Glass’ delta effect size is calculated as the standardised difference between the means of the control and treatment groups, over the standard deviation of the control group. We calculated the effect size for each behavioural trait for larvae, adult females and adult males.

## Results

### Starved larvae exhibit shorter PCI duration and higher activity

There was a significant interaction effect of treatment and the time point on PCI duration (*χ_1_^2^* = 10.78, *p* = 0.001, Figure 1a), such that starved larvae had a shorter PCI duration than control larvae at 3 h and 4 h of treatment. There was no significant effect of body mass (*χ_1_^2^* = 1.63, *p* = 0.201).

**Figure 1.**
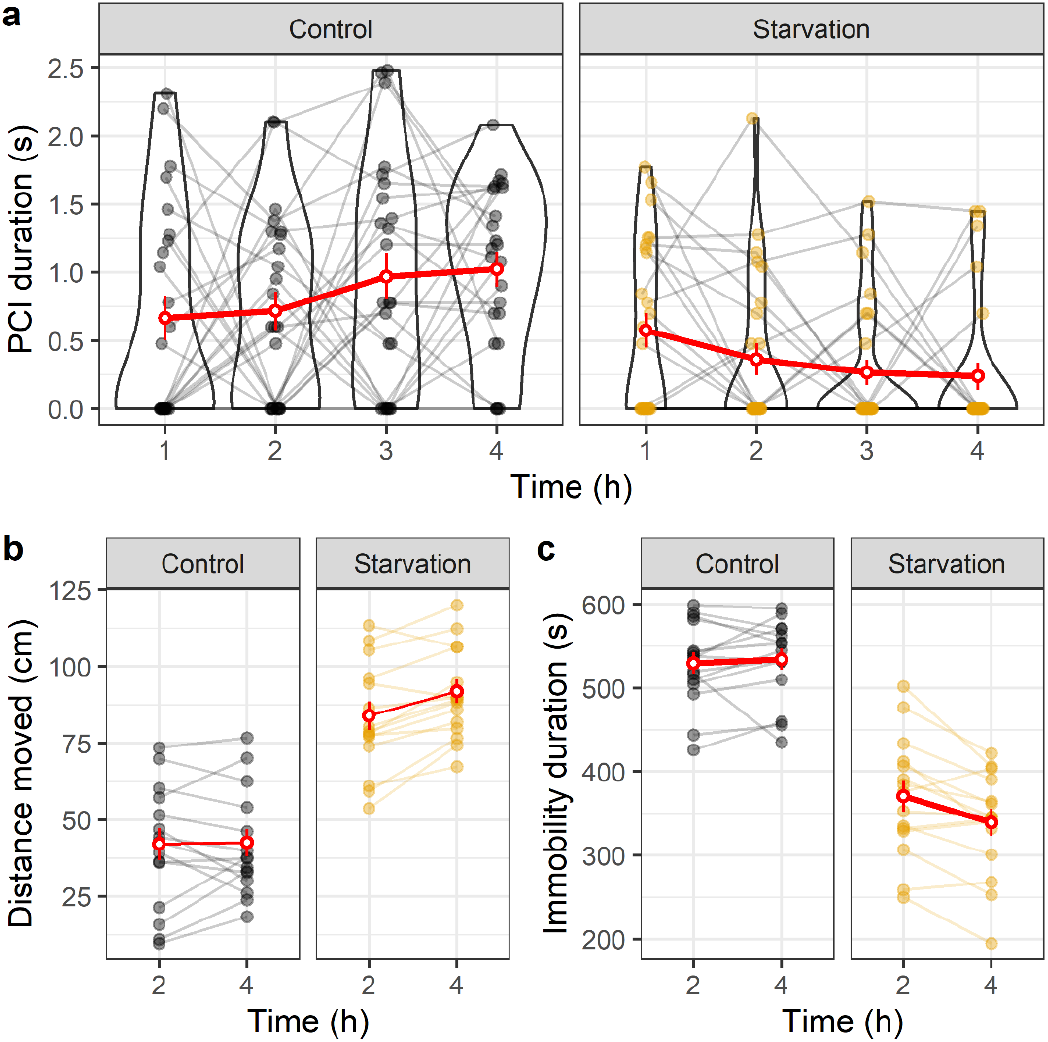
Effect of treatment (Control or Starvation) on (a) PCI duration (s), (b) distance moved (cm) and (c) immobility duration (s) over time (h) in *Athalia rosae* larvae. Red circles and error bars represent means and standard errors. For PCI duration, log-transformed raw data is plotted in transparent colours and the data distribution is visualised through violin contours. To avoid infinite values (for data point values that were zero), we added one to each data point before log-transformation (see text for details). The grey lines or coloured lines connect data over time from each larva.

There was a significant interaction effect of treatment and the time point on distance moved (*χ_1_^2^* = 4.71, *p* = 0.029, Figure 1b), but no significant effect of body mass (*χ_1_^2^* = 2.31, *p* = 0.127). Starved larvae moved significantly longer distances than control larvae. Additionally, the distance moved increased with time for larvae of the starvation treatment but not for control larvae (Supplement S2a). Similar to the distance moved, there was a significant interaction effect of treatment and the time point on immobility duration (*χ_1_^2^* = 9.77, *p* = 0.001, Figure 1c), but also a significantly positive effect of body mass (*χ_1_^2^* = 4.03, *p* = 0.044). Control larvae were immobile for a significantly longer duration than starved larvae. Moreover, the immobility duration decreased with time for larvae of the starvation treatment but not for control larvae (Supplement S2b).

### Starved adults exhibit similar PCI duration but lower activity

There was no significant effect of body mass, sex, treatment, time or the interaction between treatment and time on PCI duration in adults (Figure 2a,d; Supplement S3a). For distance moved, there was a significant effect of treatment (*χ_1_^2^* = 9.43, *p* = 0.002) and sex (*χ_1_^2^* = 5.98, *p* = 0.014), with starved individuals and males moving a shorter distance (Figure 2b,e; Supplement S3b). Similarly, there was a significant effect of treatment (*χ_1_^2^* = 9.51, *p* = 0.002) and sex (*χ_1_^2^* = 10.15, *p* = 0.001) on immobility duration, with starved individuals and males being immobile for longer durations (Figure 2c,f; Supplement S3c).

**Figure 2.**
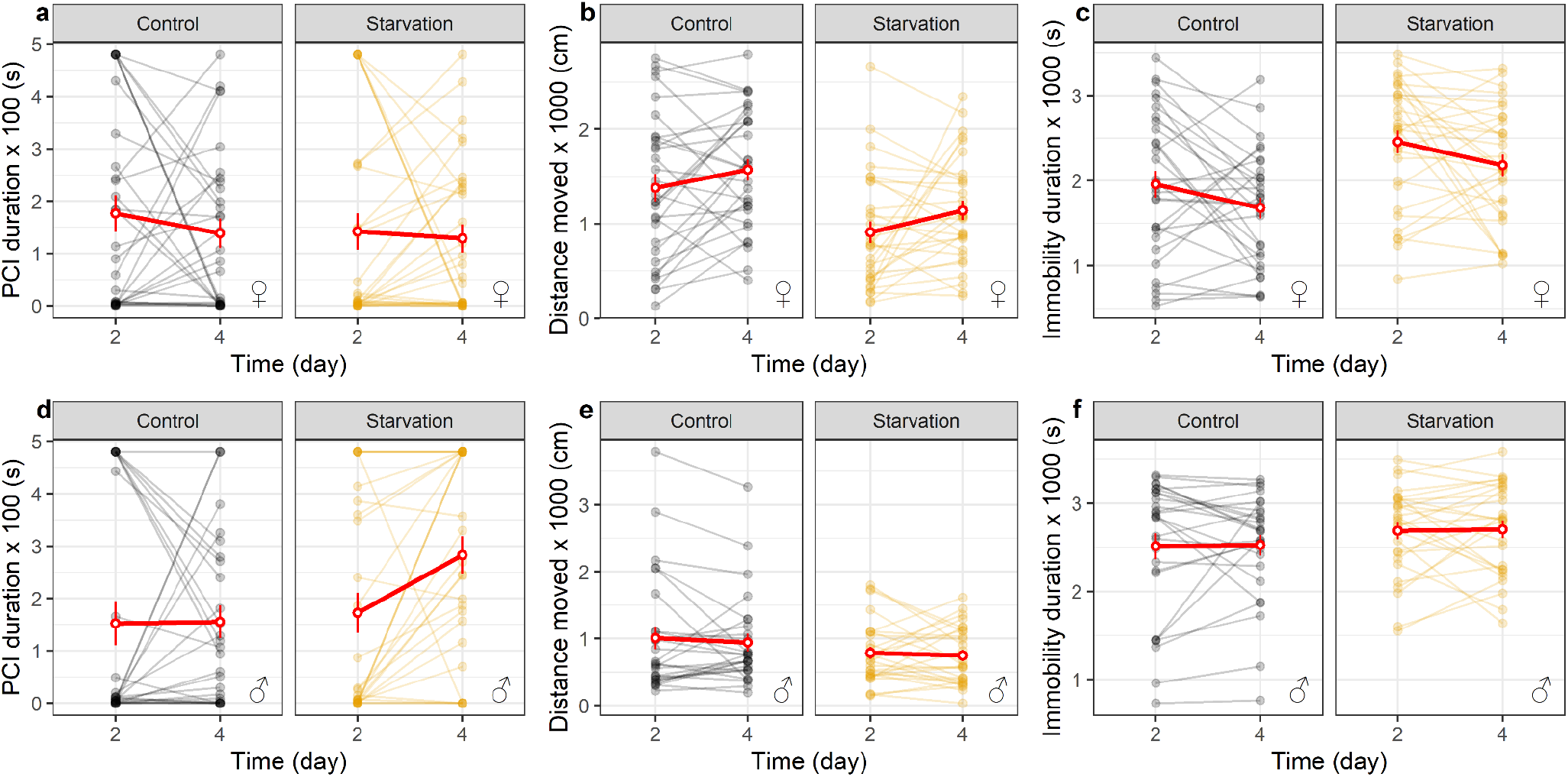
Effect of treatment (Control or Starvation) on (a) PCI duration (s), (b) distance moved (cm) and (c) immobility duration (s) over time (day) in *Athalia rosae* female (top row) and male (bottom row) adults. Red circles and error bars represent means and standard errors. The raw data are plotted in transparent colours and the lines connect data over time from each adult.

### No significant effect of starvation on repeatability estimates

Significant repeatability estimates were found for nearly all behavioural traits except for PCI duration in adult females and control treatment larvae (Table 1). While there was a trend for higher repeatability estimates for individuals of the starvation treatment for all behavioural traits of larvae and for PCI in adult males and females, this was not statistically significant (*p* > 0.05, Table 1). In contrast, for distance moved and immobility duration, control treatment individuals had a higher repeatability in adults for both sexes but this was not statistically different from adults of the starvation treatment (*p* > 0.05, Table 1).

**Table 1.** Repeatability estimates (R) with corresponding 95% confidence intervals (CI) in brackets for behavioural traits for larvae, adult males and adult females of *Athalia rosae*. Significant (*p* <0.05) repeatability estimates are highlighted in bold. The ‘*p*’ column reports the asymptotic two-tailed *p*-value for the difference in repeatability estimates between individuals of the control and starvation treatment for each behavioural trait (see text).

| Life-stage | PCI |  |  | Distance moved |  |  | Immobility duration |  |  |
| --- | --- | --- | --- | --- | --- | --- | --- | --- | --- |
| | Control | Starvation | $p$ | Control | Starvation | $p$ | Control | Starvation | $p$ |
| Larvae | 0.122 | **0.3** | 0.23 | **0.829** | **0.895** | 0.50 | **0.772** | **0.856** | 0.54 |
|  | [0, 0.317] | [0.07, 0.492] |  | [0.599, 0.945] | [0.745, 0.968] |  | [0.474, 0.921] | [0.648, 0.953] |  |
| Adult female | 0.139 | 0.21 | 0.78 | **0.591** | **0.35** | 0.24 | **0.47** | **0.417** | 0.77 |
|  | [0, 0.452] | [0, 0.537] |  | [0.335, 0.786] | [0, 0.63] |  | [0.175, 0.698] | [0.072, 0.694] |  |
| Adult male | **0.328** | **0.494** | 0.47 | **0.799** | **0.518** | 0.06 | **0.715** | **0.584** | 0.42 |
|  | [0, 0.63] | [0.172, 0.723] |  | [0.608, 0.903] | [0.189, 0.759] |  | [0.499, 0.868] | [0.258, 0.794] |  |

### Starvation has larger effect sizes during the larval than the adult life-stage

For all behavioural traits, starvation had a larger effect during the larval stage than during the adult stage in our experiments as evident by the larger Glass’ delta effect size (Figure 3a). Moreover, the direction of effect changed, as starvation during the larval stage led to more active individuals, while starvation during the adult stage generally led to less active individuals.

**Figure 3.**
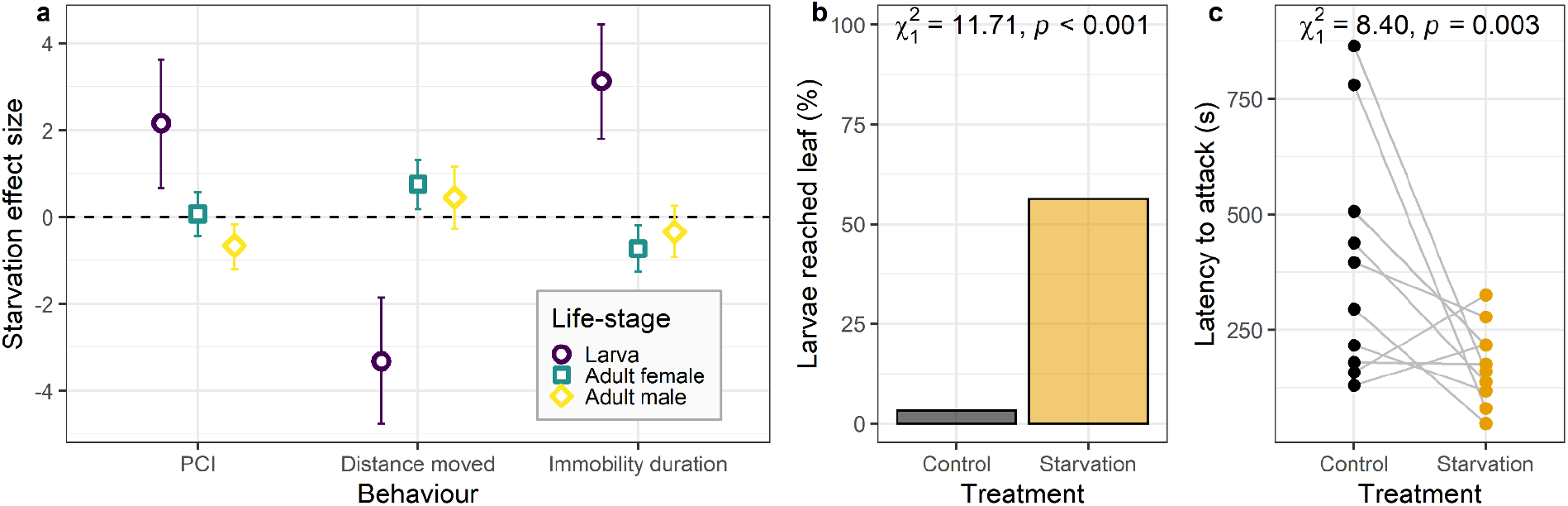
(a) Standardised Glass’ delta effect sizes for comparing starvation effect during larval and adult stage in *Athalia rosae*. For adult stage, the effect sizes were calculated separately for females and males. The error bars represent 95% confidence intervals of the effect sizes. (b) Effect of starvation on percentage of larvae that reached leaf in microcosm experiment (n = 32 replicates for each treatment). (c) Effect of starvation on latency to attack (s) by a mantid. The grey lines connect data from each mantid, tested in different order (n = 10 mantids).

### Starved larvae locate food more effectively than control larvae

Treatment had a significant effect on the probability of larvae reaching leaves (*χ_1_^2^* = 11.71, *p* < 0.001), with eighteen (56%) starved larvae reaching leaves at the end of the trials compared to only one (3%) control larva (Figure 3b, also see Supplement S4).

### Shorter latency to attack by predator for starved larvae than control larvae

Besides one mantid that refrained to launch any attack, all larvae were attacked by mantid predators and showed easy bleeding (Supplement S5a,b,c). The latency to attack by a mantid was significantly affected by larval treatment (*χ_1_^2^* = 8.40, *p* = 0.003) but not by larval body mass (*χ_1_^2^* = 0.06, *p* = 0.798), with starved larvae being attacked sooner than control larvae (Figure 3c).

## Discussion

Effect of state variables, such as body condition, on behavioural traits of an individual has been widely documented (Scharf, 2016; Rádai *et al*., 2022), but empirical evidence for how such effects may vary with life-stage is sparse. In our study we used starvation to manipulate the body condition of larvae and adults in *A. rosae*. Starved larvae showed a shorter PCI duration and higher activity levels than control larvae. This is in line with the ’asset protection principle’, which predicts that individuals in poor-condition exhibit higher risk-taking behaviour and/or activity than individuals in good condition (Catano *et al*., 2016; Kuczynski *et al*., 2016; Naman *et al*., 2019), based on their needs to reach a better state (Barclay *et al*., 2018). A low PCI duration and higher activity levels may be adaptive and allow starved larvae to disperse and locate food more rapidly. In line with this assumption, our microcosm experiment showed that starved larvae dispersed more and located food more rapidly than control larvae (also see Supplement S4a,b). Starvation usually leads to an increase in dispersal, when the benefits of foraging are higher than potential costs of exploring (Scharf, 2016). For example, a recent study with multiple nematode species and a natural nematode community showed that availability of food is an important driver of dispersal, with dispersal of nematodes increasing when food availability was limited (Kreuzinger-Janik *et al*., 2022).

However, rapid food location and increased foraging can also pose costs, such as higher predation risk (Scharf, 2016). In line with this, mantids had a shorter latency to attack starved larvae than control larvae. A faster attack could occur if the starved larvae either came more often in close contact with the mantid and/or were noticed sooner by the mantid, potentially due to their higher activity, than the control larvae. Similarly, in the Texas ratsnake, *Elaphe obsolete*, increased activity and movement have been found to be associated with higher mortality in both sexes (Sperry & Weatherhead, 2009). While we did not allow the mantid to consume the larvae, the mantids may reject the larvae also in the long term due to the easy bleeding which was evoked by all larvae in response to the initial attack (Supplement S5a,b,c). The easy bleeding brought the mantids in direct contact with glucosinolates that are sequestered from the host plants in the haemolymph (Müller *et al*., 2001) and have been shown to act deterrent on other predators (Müller *et al*., 2002; Müller & Brakefield, 2003). However, we cannot exclude that also other compounds than the glucosinolates or just the stickiness of the haemolymph may have acted deterrent on the mantids in the present experiment. Interestingly, another mantid species, *Hierodula patellifera*, has been shown to consume adults of *A. rosae* (Singh *et al*., 2022). Adults do not show the easy bleeding defence, but also contain glucosinolates in their haemolymph (Müller *et al*., 2001). However, when adults had taken up clerodanoids from another plant species, they were rejected by the mantids after an initial attack (Singh *et al*., 2022). Thus, the different life-stages of *A. rosae* seem to have different chemical mechanisms in place as protection against mantids that probably need to be presented on the outside of the prey to act effectively.

In contrast to larvae, starved *A. rosae* adults exhibited lower activity levels than control treatment adults. Lower activity in starved individuals is in agreement with the ‘state-dependent safety hypothesis’, which assumes that food-deprived individuals may be less active and less prone to take risks as the costs of risky behaviour are higher than for individuals in good conditions (Moran *et al*., 2021). Given their state, lower activity levels may allow starved individuals to conserve energy for other behaviours, such as mating. Such decreased activity levels in response to starvation have been shown in some insects like the European earwig (Weiß *et al*., 2014), two-spotted spider mite (Le Goff *et al*., 2012) and speckled cockroach (Reynierse *et al*., 1972), although such patterns do not seem to be particularly widespread (Scharf, 2016). Moreover, females were significantly more active than males. An equivalent sex-specific activity patter has been found in the leaf beetle *Phaedon cochleariae* (Müller & Müller, 2015). Such higher activity in females has been suggested to be related to the need to find oviposition sites (Müller & Müller, 2015). Sex-specific differences in behaviour have been shown in multiple species (Tarka *et al*., 2018), though the direction can vary between species and also depend on the behaviour in consideration. Some *A. rosae* adults exhibited PCI (Figure 2a,d), although there was no significant effect of any variable on PCI duration. It is possible that adults may rely on other anti-predator strategies, such as chemical defences using clerodanoids (Singh *et al*., 2022), rather than exclusively on PCI. Moreover, adults can also simply fly away, while larvae are less mobile. To our knowledge, thanatosis behaviour has previously been described in adults of only one other sawfly species, *Perreyia flavipes* (Neves & Pie, 2018).

While most repeatability estimates were significant for the behavioural traits in both life-stages of *A. rosae* in the present study, we did not find any significant effect of starvation on repeatability estimates. In the harvestman *Mischonyx cuspidatus*, individuals that were sated showed a higher repeatability in boldness levels compared to individuals from food-deprived conditions, using death-feigning duration as a proxy for boldness (Segovia *et al*., 2019). In contrast, in the black widow spider, *Latrodectus hesperus*, individuals reared on restricted diet showed repeatability for all examined behavioural traits, while individuals on an ad-lib diet exhibited repeatability only for aggression (DiRienzo & Montiglio, 2016). Thus, food deprivation can alter the repeatability of behavioural traits, although the effect can depend on the trait and species under consideration. We saw contrasting trends of higher repeatability for starved individuals compared to control individuals during the larval stage, and vice-versa in the adult stage although these differences in repeatability estimates between treatments were not significant. Future studies should examine if behavioural repeatability is impacted differently by stresses depending on the life-stage at which it is experienced.

Our results showed a large and opposite effect of starvation on behavioural traits of larvae compared to adults, with starved larvae showing a decreased PCI duration and increased activity levels while starved adults showed decreased activity levels. In contrast, in two hemimetabolous tick species the duration of PCI decreased with starvation across life stages (Oyen *et al*., 2021). Similar to our results, a recent meta-analysis revealed that the effect of poor nutrition on risk-taking behaviour could be life-stage dependent, with juveniles showing stronger responses and higher risk-taking behaviour in low-nutrition conditions while the effect on adults was less unequivocal (Moran *et al*., 2021). Life stage-specific effects on behaviour have been documented in previous studies (Polverino *et al*., 2016; Rádai *et al*., 2022), although few studies have documented such effects in holometabolous insects which occupy distinct niches in the juvenile and adult stage. The need to adjust to conditions of lower food availability may lead to individualised niche conformance (as defined in Müller *et al*., 2020).

In holometabolous insects, the uptake of food during the larval stage determines adult body mass and may also primarily determine the energy reserves available. Thus, intensive foraging allows larvae to gain body mass and reach the reproductive stage rapidly. For starved larvae, increasing activity levels may allow them to locate food even though it may be risky (Scharf, 2016). In contrast, for starved adults’ higher activity levels may deplete their already limited energy reserves and hence, reduce their opportunities for energy-costly tasks like mating and ovipositing. Moreover, individuals in larval stages or juveniles may need to invest more in foraging as their mass-specific metabolic requirements may be higher (Brown & Braithwaite, 2004). In our experiment, activity levels of larvae may have been measured under more realistic conditions than that of the adults which can fly, although we compared between control and starved treatments within each life-stage and not across life-stages. Our results demonstrate that depending on the life-stage of an individual the state-dependency of behaviour can vary and even show contrasting effects. Moreover, the adaptive value of behavioural strategies appears to be life-stage-specific for foraging, but can also be costly in terms of predation.

## Acknowledgements

This study was funded by the German Research Foundation (DFG) as part of the SFB TRR 212 (NC^3^), project number 396777467 (granted to CM). We would like to thank Karin Djendouci and Sophie Lee Beyer for their help in conducting the experiment.

## Author Contributions

Conceptualization and experimental design: PS, CM; data collection: JW for larval PCI behaviour, PS for other experiments; data validation and analysis: HS and PS for repeatability estimates and their statistical analysis, PS for other experiments; data visualization: PS; writing original draft: PS, CM; reviewing and editing: PS, HS, CM; funding acquisition: CM.

## Ethical note

This study complies with the current laws of Germany on the use of invertebrates in research and adheres to the ASAB/ABS Guidelines for the Use of Animals in Research.

## Data availability

The data set is available on the repository Zenodo https://doi.org/10.5281/zenodo.7437089.

